# ALT in Pediatric High-Grade Gliomas Can Occur without *ATRX* Mutation and is Enriched in Patients with Pathogenic Germline MMR Variants

**DOI:** 10.1101/2022.08.05.502870

**Authors:** Jennifer L. Stundon, Heba Ijaz, Krutika S. Gaonkar, Rebecca S. Kaufman, Run Jin, Anastasios Karras, Zalman Vaksman, Jung Kim, Ryan J. Corbett, Matthew R. Lueder, Daniel P. Miller, Yiran Guo, Mariarita Santi, Marilyn Li, Gonzalo Lopez, Phillip B. Storm, Adam C. Resnick, Angela J. Waanders, Suzanne P. MacFarland, Douglas R. Stewart, Sharon J. Diskin, Jo Lynne Rokita, Kristina A. Cole

## Abstract

**Background:** To achieve replicative immortality, most cancers develop a telomere maintenance mechanism, such as reactivation of telomerase or alternative lengthening of telomeres (ALT). There are limited data on the prevalence and clinical significance of ALT in pediatric brain tumors, and ALT-directed therapy is not available.

**Methods:** We performed C-circle analysis (CCA) on 579 pediatric brain tumors that had corresponding tumor/normal whole genome sequencing through the Open Pediatric Brain Tumor Atlas (OpenPBTA). We detected ALT in 6.9% (n=40/579) of these tumors and completed additional validation by ultrabright telomeric foci *in situ* on a subset of these tumors. We used CCA to validate *TelomereHunter* for computational prediction of ALT status and focus subsequent analyses on pediatric high-grade glioma (pHGG) Finally, we examined whether ALT is associated with recurrent somatic or germline alterations.

**Results:** ALT is common in pHGG (n=24/63, 38.1%), but occurs infrequently in other pediatric brain tumors (<3%). Somatic *ATRX* mutations occur in 50% of ALT+ pHGG and in 30% of ALT-pHGG. Rare pathogenic germline variants in mismatch repair (MMR) genes are significantly associated with an increased occurrence of ALT. Conclusions: We demonstrate that *ATRX* is mutated in only a subset of ALT+ pHGG, suggesting other mechanisms of *ATRX* loss of function or alterations in other genes may be associated with the development of ALT in these patients. We show that germline variants in MMR are associated with development of ALT in patients with pHGG.

**Key Points:** ATRX alterations are frequent, but not required, for an ALT phenotype in pHGGs

pHGG patients with germline mismatch repair variants have higher rate of ALT + tumors *TelomereHunter* is validated to predict ALT in pHGGs

**Importance of the Study:** We performed orthogonal molecular and computational analyses to detect the presence of alternative lengthening of telomeres in a highly characterized cohort of pediatric brain tumors. We demonstrate that many pHGG utilize ALT without a mutation in ATRX, suggesting either loss of function of ATRX via an alternative mechanism or an alternate means of development of ALT. We show that germline variants in MMR genes are significantly associated with ALT in pHGG. Our work adds to the biological understanding of the development of ALT and provides an approach to stratify patients who may benefit from future ALT-directed therapies in this patient population.

## Introduction

As human cells divide, telomeres become progressively shorter, leading to senescence^1^. Cancer must overcome this barrier to achieve replicative immortality^2^. In most cancer, telomeres are maintained via telomerase reactivation, however, in 10-15% of cancers, telomeres are lengthened by the recombination-based mechanism, alternative lengthening of telomeres (ALT)^3^. ALT occurs most frequently in mesenchymal tumors and has been shown to occur frequently in pHGG, neuroblastoma, osteosarcoma, as well as adult low-grade glioma and pancreatic neuroendocrine tumors^4,5^. Additionally, ALT has been observed in some medulloblastomas and primitive neuroectodermal tumors^6^. ALT is rarely found in epithelial malignancies, likely because telomerase expression is less tightly regulated in epithelial cells^7^. While ALT has been shown to frequently occur in pediatric pHGG, previous studies have not integrated associated tumor and normal whole genomes with clinical data for these patients^8^.

ALT+ telomeres utilize homologous recombination to maintain telomere length and have a high level of telomeric DNA damage secondary to replication stress^9^. ALT can be identified by measuring the presence of ultra-bright telomeric foci, extra-chromosomal C-circles, or ALT-associated pro-myelocytic bodies^3,10,11^. Additionally, computational methods can identify and count telomere repeats from sequencing data^12–15^, though previous studies have utilized limited pediatric brain tumors^16^. Collectively, these techniques measure the presence of DNA damage, replication stress, and altered telomeric content that are characteristic of ALT telomeres. The use of ALT as a telomere maintenance mechanism (TMM) has been associated with poor outcome in some cancers, such as neuroblastoma, though in other cancers including adult glioblastoma, patients with ALT+ tumors have a better prognosis compared to patients with telomerase positive cancer^17,18^. It remains unclear whether there is a prognostic difference among ALT+ pHGG patients compared to those with ALT-tumors.

Loss of function mutations in the *ATRX* chromatin remodeling gene are strongly correlated with ALT positivity and one known role of *ATRX* is to inhibit ALT^19–21^. A recent study in pediatric neuroblastoma showed that reduced protein abundance of *ATRX*, was only observed in 55% of ALT+ patients^17^ and it has been shown that *ATRX* mutations are not required for activation of ALT in adult pancreatic neuroendocrine tumors and melanoma^22,23^.

It is not known whether germline variants in DNA repair genes are associated with the development of ALT. Recent literature suggests that loss of MMR function may have an important role in ALT activity in human cancer cell lines^24,25^, though this association has not yet been demonstrated in primary human tumors. Additionally, it has been shown that loss of MMR function in yeast and mice is associated with telomerase-independent telomere lengthening and improved organismal survival and fitness^26,27^, which supports a role for loss of the MMR pathway in promoting the development of ALT. Cancer predisposition syndromes such as constitutional MMR deficiency (CMMRD) and Lynch Syndrome (LS), as well as acquired somatic MMR gene alterations, are major mechanisms of MMR pathway loss of function. CMMRD is a rare, aggressive predisposition syndrome resulting from biallelic pathogenic germline variants in the MMR genes *PMS2* (60%), *MSH6* (20-30%), *MLH1*/*MLH2* (10-20%). LS has an autosomal dominant mode of inheritance and is caused by monoallelic pathogenic germline variants in the same MMR genes: *MSH2/MLH1* (80%), *MSH6* (13%), and *PMS2* (6%)^28^.

Here, we assess the frequency of ALT, as well as clinical and molecular phenotypes associated with ALT, in a large cohort of pediatric brain tumors from the OpenPBTA, with detailed investigation of pHGGs^29–31^. We validate the use of the computational algorithm, *TelomereHunter*^*16*^, to predict ALT status. We demonstrate that *ATRX* is only mutated in a subset of ALT+ pHGG patients, and that presence of pathogenic germline variants in MMR genes is strongly associated with the development of ALT. This is important, as germline variants in MMR genes are observed in roughly 6% o pHGG tumors^32^, a common tumor observed in patients with LS and CMMRD^33^. By demonstrating that MMR variants are associated with ALT and that *ATRX* is only mutated in a subset of ALT+ pHGG tumors, we add to our understanding of the key molecular changes that are associated with ALT. Developing a greater understanding of the molecular drivers of ALT will be critical to the creation of ALT-directed therapy.

## Methods

### Pediatric brain tumor data and genomic analyses

#### TelomereHunter

Paired tumor and normal WGS BAMs (N = 940) from previously sequenced pediatric brain tumors were obtained by data access request to the Children’s Brain Tumor Network (CBTN)^34^. The BAMs were used as paired inputs to *TelomereHunter*^16^, which was run using default parameters to estimate telomere content. The ratio of telomere content in a tumor compared to its normal was calculated and used for all downstream analyses. Using the C-circle assay readout as a positive or negative ALT phenotype (N = 579 samples), we used the R package *cutpointr*^35^ to determine a telomere ratio cutoff to assign samples as ALT+ while also estimating accuracy, sensitivity, and precision. To further validate receiver operating characteristic (ROC) curves, we randomly shuffled telomere ratio scores and plotted the ROC for shuffled scores.

#### Assessment of germline variant pathogenicity

Germline variants in genes included in the KEGG MMR gene set, plus *POLE* (**Table S3**) were first annotated using SnpEff v4.3t, ANNOVAR (06-07-2020. Variant with read-depth ≥ 15, variant allele fraction ≥ 0.20, and observed in < 0.1% across each population in the public control databases non-TCGA ExAC (exonic) or gnomAD 2.1.1 (non-exonic, splicing) were considered for further study. We retained variants annotated as Pathogenic/Likely Pathogenic (P/LP) in ClinVar (05-07-2022) or InterVar v2.2.2. All Pathogenic/Likely Pathogenic (P/LP) calls were manually reviewed by an interdisciplinary team, including clinicians and genetic counselors.

### C-Circle Analysis

Tumor DNA was obtained from the Children’s Brain Tumor Network^34^ from individuals consented on the institutional review board approved CBTN protocol. C-circle analysis (CCA), which is a highly accurate read-out of ALT^36^, was performed on tumor samples and controls as described previously^10^. Quantification of positivity was performed as described previously^37^.

### Tissue Microarray

Full methods detailing creation of tissue microarray, ATRX IHC, H3K28me3 IHC, H3K28M IHC, and UBTF analysis are included in supplementary methods.

### Statistical tests

Fisher exact tests were used to determine statistical significance for categorical variables. Mann-Whitney U testing was used to compare populations of two groups.

## Results

### Frequency of ALT in pediatric brain tumors

To characterize ALT in pediatric brain tumors, we performed CCA on 579 tumors from unique patients which had corresponding sequencing data available in the OpenPBTA (**Figure 1A**). We found a low frequency of ALT in ATRT (N = 2/23, 8.7%), ependymoma (N =3/64, 4.7%), ganglioglioma (N = 1/41, 2.5%), and medulloblastoma (N = 4/86, 4.7%). ALT has not been previously described in ATRT or ependymoma^6^. In contrast, we confirmed that 38.1% of pHGG (N = 24/63) have ALT. This is concordant with previous studies reporting that approximately 40% of pediatric pHGG utilize ALT^5,6,8^. We orthogonally validated the CCA by measuring ultra-bright telomeric foci (UBTF) via telomere FISH in 28 pHGGs. We show that 100% of tumors which were CCA positive were also positive for UBTF (N = 8), and only 2 (10%) of the remaining 20 tumors which were CCA negative were positive by UBTF (**Supplementary Table 1, Figure 2**). Since the majority of ALT tumors in our study were pHGG, we focused subsequent analyses on the pHGG cohort (N = 85: *N = 63 with CCA + UBTF, N = 20 with WGS only, N = 2 with WGS+UBTF*, denoted as the “primary analysis” cohort) shown in **Figure 1B. Of these, 41 were H3 K28-altered, 40 were H3 wildtype, and 4 were H3 G35-mutant (Figure 1C)**.

**Figure 1.**
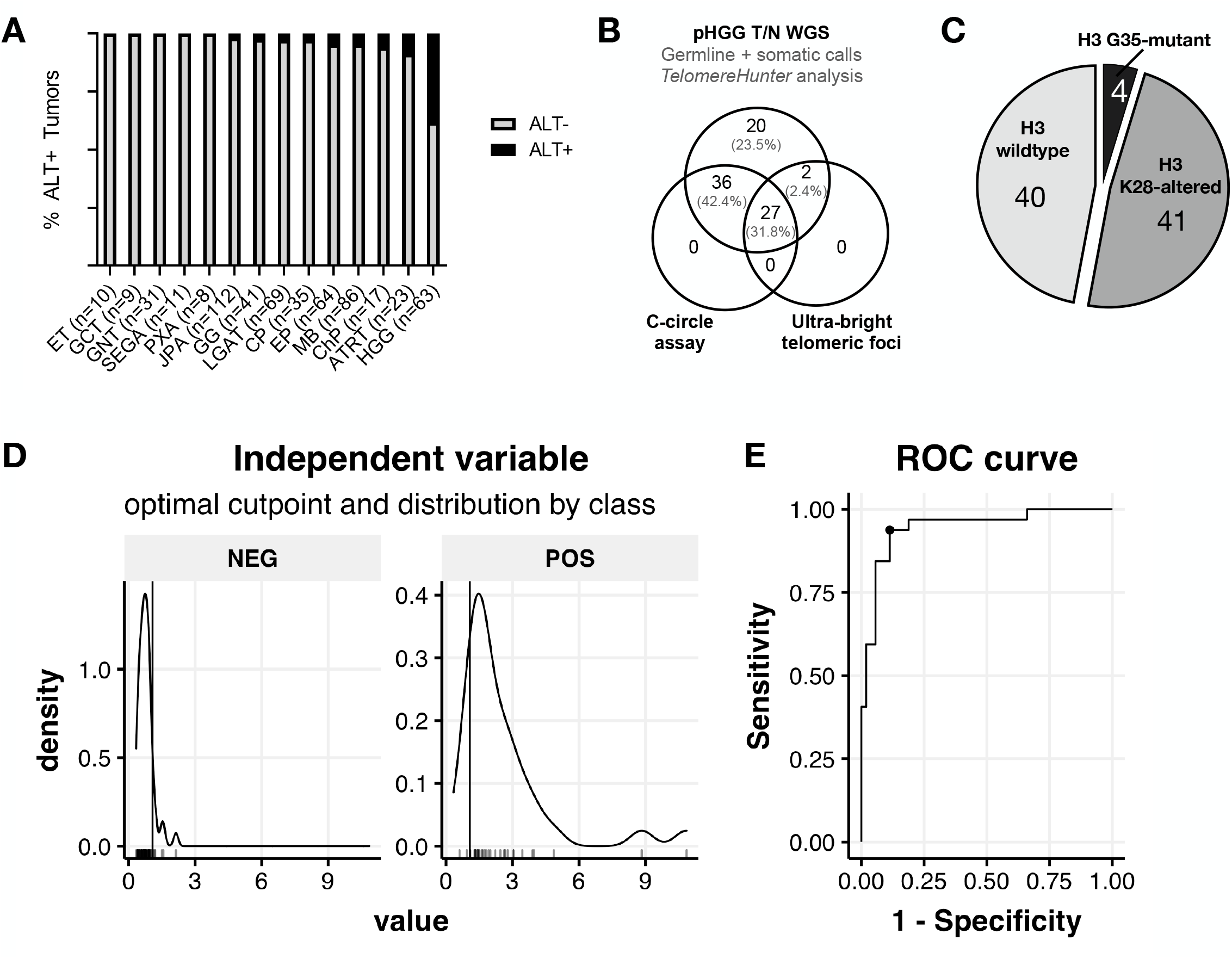
ALT is more prevalent in pediatric pHGGs than other CNS tumors and can be computationally determined. (**A**) C-circle analysis was completed for PBTA primary tumor samples from unique pediatric patients (N = 579). Tumor abbreviations: ET= Embryonal tumors, including CNS embryonal tumors and embryonal tumors with multilayered rosettes, GCT = germ cell tumor, GNT = glioneuronal tumor, LGAT = other low grade astrocytic tumor, JPA = juvenile pilocytic astrocytoma, PXA = Pleomorphic xanthoastrocytoma, SEGA = Subependymal giant cell astrocytoma, GG = ganglioglioma, CP = craniopharyngioma, EP= ependymoma, MB = medulloblastoma, ChP= choroid plexus tumor, ATRT = atypical teratoid rhabdoid tumor, HGG = high grade glioma. Benign lesions, non-primary brain tumors, tumors from patients >21 years at time of diagnosis, tumors with fewer than five samples per disease type and duplicate samples for a single patient were excluded from this analysis. pHGG represents 10.8% of tumors analyzed, but 60% of ALT+ tumors. (**B**) Representation of the pHGG subset (N = 85) on which paired tumor/normal WGS was performed. There was sufficient DNA available to perform C-circle assay on 63 samples and ultra-bright telomeric foci analysis was performed on 24 samples. (**C**) Using the C-circle assay data as the “truth” set for an ALT phenotype, we used the R package *cutpointr* to determine the optimal tumor/normal telomere content ratio cutpoint for determining ALT +/-status. Shown are density plots for ALT + or - pHGG samples at a cutpoint ratio of 1.0679 (x-intercept). (**E**) This cutpoint enabled a 90.59% accuracy, 93.75% sensitivity and 88.68% specificity, shown with the receiver operating characteristic (ROC).

**Figure 2.**
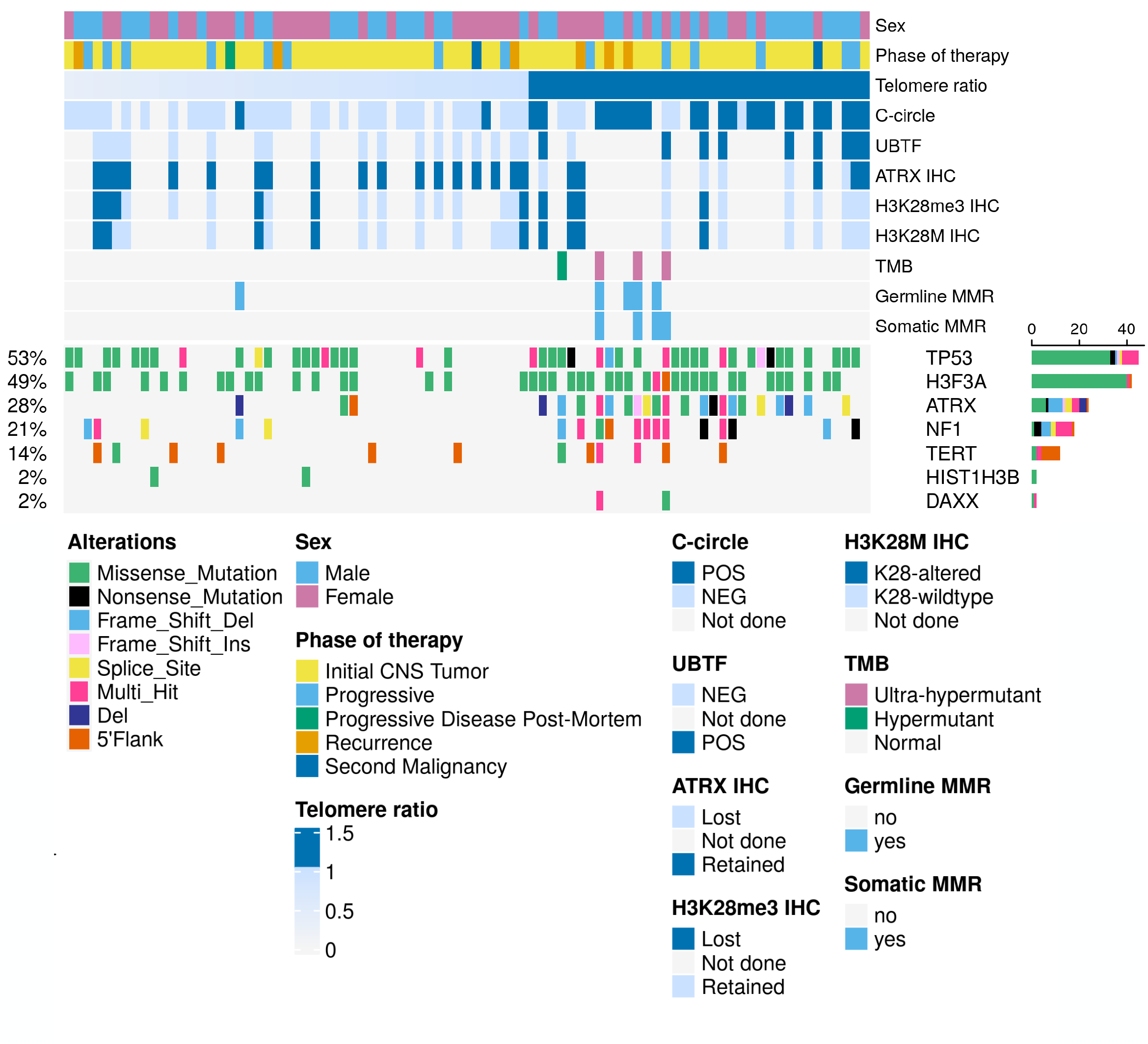
Molecular phenotypes and genomic alterations of pediatric pHGG tumors by ALT status. Annotations for sex (estimated from germline WGS), tumor phase of therapy, *TelomereHunter* telomere ratio (tumor v. normal), C-circle assay, ultrabright telomeric foci assay, and immunohistochemistry for ATRX, H3 K28M and H3 K28me3 are shown. TMB is annotated for hypermutant (100 Mut/Mb > TMB >= 10 Mut/Mb) and ultra-hypermutant (TMB >= 100 Mut/Mb) tumors. Positivity for variants in germline and/or mutations in somatic mismatch repair (MMR) genes (listed in **Table S3**) is annotated above the individual somatic mutations in *TP53, H3F3A, ATRX, NF1, TERT, HIST1H3B*, and *DAXX*.

### TelomereHunter accurately predicts an ALT phenotype in pediatric pHGG

Computational methods of identification of ALT can provide rapid prediction of ALT in some patient tumors and may become important clinical tools as ALT-directed therapies are developed^16^. To determine whether a computational method of ALT identification could be utilized to detect ALT in pediatric brain cancer, we used *TelomereHunter*^*16*^ to estimate telomere content from paired tumor/normal WGS (N = 940) and calculated the tumor/normal telomere content ratio (**Figure 1 -D-E** and **Supplementary Figure S1**). Using the molecular readout for pHGG patients on which CCA was performed (N = 63), we validated the utility of *TelomereHunter* to accurately stratify pHGG tumors by ALT status. We determined that a tumor/normal telomere content ratio of >1.0679 could identify ALT in pHGG (ROC = 0.95), achieving 90.59% accuracy, 93.75% sensitivity, and 88.68% specificity (**Figure 1D** and **1E**). This demonstrates that the use of *TelomereHunter*, which can be performed on any patient tumor with paired normal and/or tumor whole genome sequencing, can identify ALT in pHGG with high accuracy. We were additionally able to identify a tumor/normal telomere content ratio of >0.9963 for non-HGG tumors (ROC = 0.66), though at a lower accuracy (76.97%), sensitivity (56.34%), and specificity (79.25%) (**Supplementary Figure S1, A-B**). This lower ROC may be the result of histology heterogeneity and/or the lower number of tumors in this group positive from CCA. However, randomized telomere content ratios (**Supplementary Figure S1C**) resulted in an expected diagonal (ROC = 0.53), suggesting the signal in both pHGG and non-HGG groups is real. We found that an ALT phenotype was predicted for tumors across all histologies except for subependymal giant cell astrocytoma (**Supplementary Figure S1D**).

To expand the cohort of pHGG patients for downstream clinical and genomic analyses, we used *TelomereHunter* to assign ALT phenotypes to 22 pHGG previously sequenced tumors without sufficient DNA for CCA to increase the pHGG cohort size to N = 85, including 53 ALT+ and 32 ALT-patients (**Table 1**).

#### Older age is associated with a higher frequency of ALT

We analyzed the clinical status of patients with pHGG with and without ALT (**Table 1**). We demonstrate that amongst all pHGG patients in our cohort, patients with ALT+ pHGG were significantly older (11.06 years vs. 7.9 years, p= 0.007, **Table 1**). This suggests that there may be inherent biologic differences in tumors of patients with older ages that contribute to the development of ALT or that older patients may be more likely to develop tumors that are more frequently associated with ALT. For example, all patients with diffuse hemispheric glioma H3 G35-mutant pHGG were ALT+, as has been previously reported^38^. The age range for these patients is 14.8-18.4 years, and this subtype of pHGG is known to be more common in adolescence and young adults ^39^. Amongst patients with diffuse midline glioma, H3 K28-altered subtype and pHGG H3-wildtype we saw no difference in rates of ALT (N= 16/40 and N= 12/41, respectively, **Table 1**). To further characterize the pHGG tumors, we performed H3K28me3 IHC on TMA samples. We show that 100% of tumorsclassified genomically as “DMG, H3 K28-altered” which were present on the TMA all had loss of H3K28me3 staining (N= 13/13) and that two tumors previously classified as H3 WT had loss of H328me3 by IHC (6%, N = 2/32). We show no difference between race or ethnicity when comparing ALT+ and ALT-patients. We further analyzed to assess differences in somatic mutations, mutational burden, and germline mutations (**Figure 2**).

#### Genomic landscape of ALT positive or negative pHGGs

Depicted in **Figure 2** is an oncoprint of 85 pediatric pHGG tumors ordered by *TelomereHunter* tumor/normal telomere content ratio. Selected clinical demographics, molecular assay results, and genomic alterations are displayed. Notably, we observed high (predicted ALT+) tumor/normal telomere content ratios in all CCA positive cases except two tumors. Likewise, all samples positive for UBTF were predicted as ALT+ by *TelomereHunter*. Additionally, we show an inverse relationship between samples positive for ATRX protein and those positive for CCA and/or UBTF, consistent with previous work^8^. This inverse relationship also extends to somatic DNA alterations in *ATRX:* samples with somatic alterations in *ATRX* generally have loss of ATRX protein. We illustrate that *ATRX* is not mutated in all ALT+ pHGG, and that mutations in *DAXX* and *TERT* were rare in our cohort. We did not find mutual exclusivity between *TERT* promoter mutations and ALT positivity, suggesting the presence of dual mechanisms of telomere maintenance in many tumors. Intriguingly, we note that half of *ATRX* WT samples (16/38, 50%) do not harbor a mutation in the *ATRX*-interacting tumor suppressor, *DAXX*. We found slight enrichment of ALT in tumors with two of the most frequently occurring somatic alterations in pediatrics pHGGs, *TP53* and *H3F3A* (p=0.027 and p=0.047, respectively). ALT was not enriched in *NF1* mutated tumors (**Supplementary Figure S2A**). Finally, we show that hypermutant and ultra-hypermutant tumors, as well as tumors with either germline or somatic MMR (**Table 2** and **Table S3**), were more likely to be ALT+ by CCA and/or *TelomereHunter*.

### Somatic *ATRX* mutations occur in 50% of ALT-positive pHGG

We sought to determine the frequency of *ATRX* mutations in our patient cohort and whether any other recurrent somatic mutations occur more frequently in ALT+ patient tumors. We did not detect rare pathogenic germline *ATRX* variants in this cohort. Somatic *ATRX* alterations were present in 50% of ALT+ pHGG patient tumors (N = 16/32, **Figure 3A, Supplementary Table 2**). *ATRX* mutations were rare in ALT-pHGG, and these were frequently variants of uncertain significance (N = 3/5). In contrast, mutations in *ATRX* in ALT+ pHGG were likely to be oncogenic (N = 13/16, **Figures 3A and Supplementary Figure S2, B-C)**. We next compared the frequency of ALT in *ATRX* WT versus *ATRX* mutant pHGG and found 76% of *ATRX* mutated pHGG (N = 16/21) are ALT+, compared to 28% (N = 16/64) of *ATRX* WT tumors (p=0.0046, **Figure 3B**).

**Figure 3.**
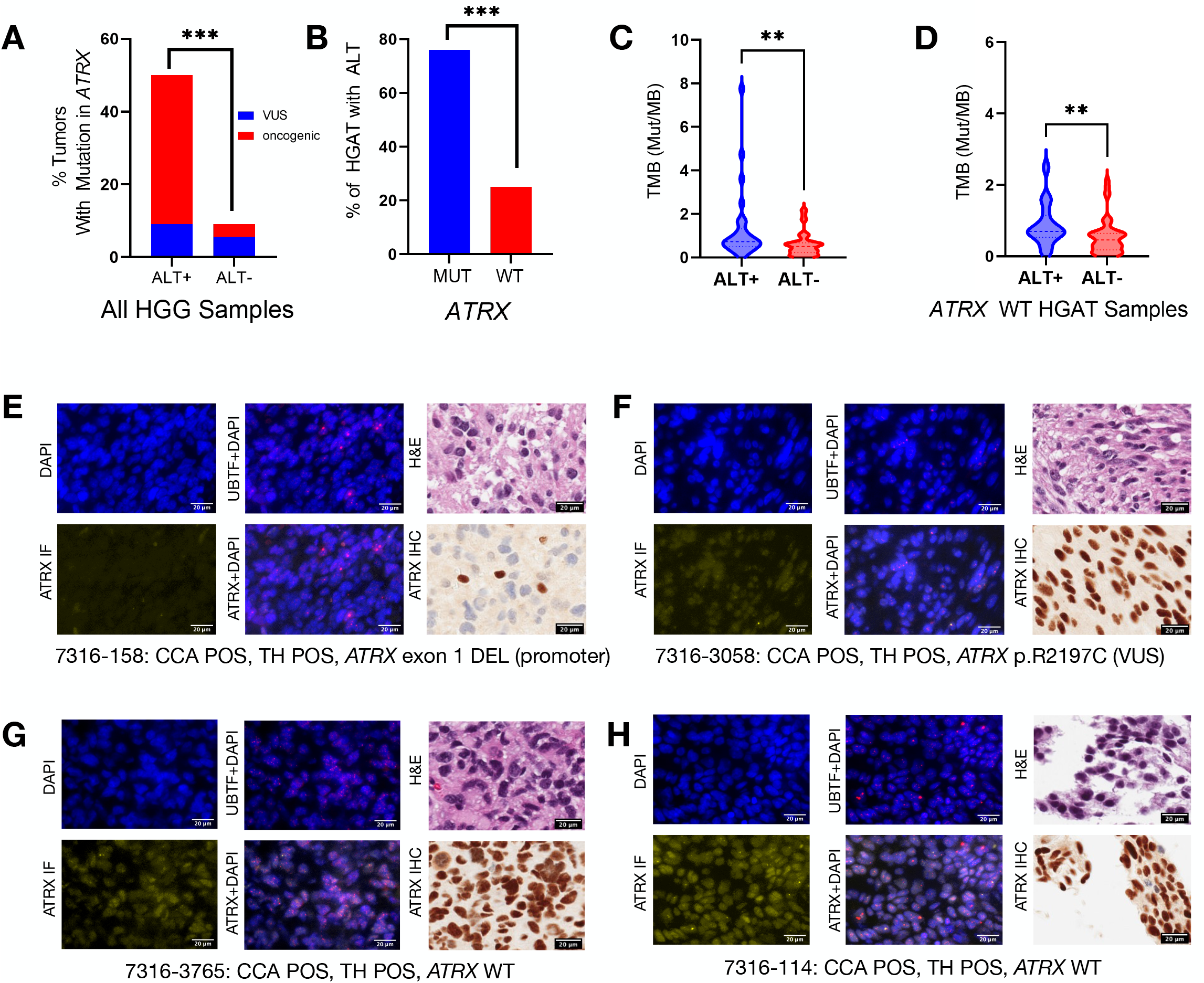
ALT+ pHGGs are significantly enriched for *ATRX* mutations and have a higher tumor mutation burden. (**A**) pHGG patients with ALT are more likely to have *ATRX* mutations (p < 0.001, N = 16/32 ALT+, N = 5/53 ALT-). *ATRX* mutations in ALT+ pHGG are more likely to be likely oncogenic mutations (N = 13/16) compared to mutations in ALT-pHGG (N = 2/5). (**B**). Mutations in *ATRX* are significantly associated with ALT (p < 0.001). (**C**) ALT+ pHGGs have a higher TMB than ALT-pHGGs (p=0.0038). (**D**) *ATRX* WT ALT+ tumors have a higher TMB compared to *ATRX* WT ALT-pHGG tumors (p=0.0098). pHGG tumors may be ALT positive with (**E**) or without (**F-H**) ATRX protein expression *in situ*. Left and middle panels: Representative images of multiplex immunofluorescence of UBTF (red), ATRX protein (yellow) or both within DAPI stained nuclei (dark blue) of ALT+ pHGG tissues from 4 patient tumors (**E**: 7316-158; **F**: 7316-3058; **G**: 7316-3765; **H**: 7316-114). Right panels: representative H&E images and ATRX IHC, noting that in E (tumor 7316-158) ATRX protein expression is absent in tumor nuclei (blue) with positive ATRX staining in non-tumor nuclei. The remaining ALT+ pHGG tumors (**F-H**) demonstrate ATRX protein staining.

To gain a greater understanding of the *ATRX* biology in our cohort, we performed ATRX immunohistochemistry (N= 30) using the same antibody and conditions as our clinical laboratory, to assess for presence of absence of the ATRX protein on the pHGG brain tumor TMA. We demonstrate that 25% of the CCA positive pHGG tested (N = 3/12) in this cohort retained ATRX protein expression, demonstrating a similar discordance between ATRX staining and ALT status as has been previously reported^40^. Representative images of ALT+ patients with ATRX protein expression retained (**Figure 3E-F**) and lost (**Figure 3G**) are depicted and full details are provided in **Supplementary Table 1**. Additionally, ATRX protein expression in *ATRX*-mutant samples is not always heterogeneous (**Supplementary Figure 3A-B**). Thus, we believe that neither *ATRX* mutation nor loss of ATRX protein expression should be used as a primary biomarker for ALT in pHGG, aligning with similar reports of ALT in neuroblastoma^17^. Notably, we show that in the pediatric brain tumor population, ALT can occur independent of somatic ATRX alterations (**Figure 3B, G-H**). Finally, we investigated the clonality of the ATRX mutations as well as their relationship with ALT status. Although there was a small trend toward higher ATRX variant allele frequencies (VAF) in ALT+ tumors (mean VAF = 0.64) compared to ALT-tumors (mean VAF = 0.51), we did not find significant differences in ATRX mutation clonality between the distributions (KS-test D = 0.35, p-value = 0.8339). This non-significant effect also remained true when accounting predicted oncogenicity of the mutation (KS-test_ALT+ onco vs. ALT+ VUS_ D = 0.54545, p-value = 0.3473 and KS-test_ALT+ onco vs. ALT-VUS_ D = 0.38636, p-value = 0.7736, Supplementary Figure 3C). We observe a positive correlation trending toward significance (Pearson’s R = 0.449, p = 0.0536) between somatic ATRX VAF and a higher TelomereHunter T/N telomere content ratio and this was not driven by tumor purity (Supplementary Figure 3D). All *ATRX*-mutant tumors in the analysis cohort harbored clonal (VAF >= 0.20) *ATRX* mutations. This perhaps suggests a requirement of *ATRX* clonal selection in *ATRX*-driven ALT, though more thorough experimental and computational analyses, including at a single cell level, should be performed to confirm this.

### Germline variants in MMR genes are associated with ALT

We were specifically interested in whether germline variants in the mismatch repair (MMR) pathway may be associated with the development of ALT, as previous work in model organisms and in a cancer cell line have suggested a relationship between MMR and ALT^24–27^. Tumors with germline variants in MMR and/or *POLE* make up approximately 8% of patients with pediatric pHGG^32^. Additionally, germline MMR variants combined with acquired somatic mutations in MMR genes or *POLE* are known to result in ultra-hypermutated patient tumors ^41^. We sought to understand whether there was a relationship between germline MMR variants and ALT in our cohort.

Using analysis of paired tumor and normal whole genome sequencing, we identified that 7% (N = 6/85) of patients with pHGG harbor heterozygous pathogenic germline variants in MMR genes (**Supplementary Tables 2 and 3**). Of the six pHGG tumors we identified with pathogenic or likely pathogenic germline MMR variants, five were ALT+, whereas 27/79 tumors without germline variants were ALT+ (p=0.02, **Table 2, Figure 2**). These results suggest that loss of function of the MMR pathway may be associated with ALT.

We reviewed the pathology reports where available and identified one additional patient in our cohort with a self-reported *PMS2* variant who was not identified in the unbiased germline variant analysis. This patient had a negative CCA but was noted to have positive UBTF and a positive *TelomereHunter* score. Two additional tumors from patients with clinically known MMR germline variants (LS and CMMRD) were analyzed on our TMA and were not part of the PBTA cohort. For both patients, UBTF was positive though only one patient had a positive CCA. TMA data for a patient with germline MMR is shown in **Figure 3F**.

Together, this demonstrates that patients with a germline variant in MMR or a clinically-diagnosed MMR disorder, such as CMMRD or LS, have a higher frequency of ALT positive pHGG tumors as compared to patients without germline variants or clinically diagnosed MMR syndromes.

### Tumor mutational burden is higher in ALT+ pHGG

Patients with germline MMR variants, particularly if biallelic, are known to have extremely high tumor mutational burden^42^, and we replicate this in our cohort (**Table 2**). We sought to explore whether ALT is associated with increased TMB among the 85 pHGG patients in the PBTA. We excluded patients with hypermutant or ultra-hypermutant status (>10 mutations/Mb or >100 mutations/Mb) and showed that ALT+ pHGG have a higher tumor mutational burden (**Figure 3C**, p=0.0022). This suggests that among patients with ALT+ pHGG, there may be tolerance for a greater level of DNA damage. Previously, somatic mutations in *ATRX* have been associated with higher TMB in pediatric pHGG, but the association between ALT and TMB has not been previously examined^43^. We therefore sought to determine whether our finding is independent of *ATRX* mutation status.We demonstrate that among *ATRX* WT tumors, there is a significant increase in tumor mutational burden in ALT+ pHGG as compared to ALT-pHGG (p=0.0098, **Figure 3D**) suggesting that *ATRX* mutation alone does not account for the increase in TMB observed in ALT+ tumors.

### ALT status alone is not predictive of overall survival for pHGG patients

We assessed the impact of ALT status on overall survival (OS), first without any covariates and then by ALT status and histone H3 subtype. We did not find a significant effect of ALT status on OS in pHGGs (p = 0.499). When stratifying by histone mutation status, there was no difference in OS in patients with H3 WT ALT + (median OS = 27.7 months) and H3 WT ALT-tumors (median OS = 29.8 months). Using additive multivariate cox regression, we report a significantly worse OS in patients with K28M tumors compared to those with H3 WT ALT-tumors, whether ALT+ (median OS = 14.4 months, HR = 3.2, p = 0.002) or ALT-(median OS = 9.1 months, HR = 4.3, p < 0.001), though we observed non-significant trends of higher OS in patients with ALT (**Supplementary Figure S4**). H3 G35-mutant pHGG (median OS = 69.9 months) were all ALT+ tumors (**Table S4**).

## Discussion

We analyzed primary patient tumors for ALT in 579 pediatric patient tumors using the gold standard CCA assay from a well-characterized pediatric brain tumor cohort. We identified ALT at low frequency in ATRT, ependymoma and medulloblastoma. We used a cohort of 85 unique patients with pHGG to analyze the clinical, demographic, and molecular differences in ALT+ and ALT-pHGG and demonstrate that *ATRX* is somatically-altered in 50% of ALT+ pHGG, ALT+ tumors have a higher mutational burden, and presence of pathogenic germline variants in MMR genes is strongly associated with the development of ALT. By correlating CCA and *TelomereHunter* output, we were able to further validate *TelomereHunter* as a reliable tool to predict ALT status in pHGG using whole genome sequencing data, which may have important clinical implications when ALT-directed therapies are available clinically.

Survival differences have been seen in other ALT+ and ALT-cancers and collecting additional samples may clarify whether survival is prolonged in any subset of pHGGs. For example, the presence of ALT extends overall survival in adult GBM^21^. It will be important to understand whether we can identify a true causal relationship in pHGG, or whether the presence of ALT is associated with other changes that confer a more favorable outcome. For example, we have demonstrated that ALT+ pHGG have a higher tumor mutational burden, and it is possible that acquisition of specific mutations may improve overall survival, or that a high tumor mutational burden creates an instability in the tumor genome that favors a longer survival. Future work will include a larger patient cohort to determine if there are other key differences in ALT+ and ALT-pHGG patients that impact survival, focusing on the K28-altered pHGG tumors.

In our primary analysis cohort, *ATRX* mutations occur in only 50% of our ALT+ pHGG patients, however, we failed to identify any other somatic mutations that may drive ALT, and *DAXX* was not mutated in any *ATRX* WT ALT+ tumors. It is possible that ALT+ tumors without *ATRX* mutations have changes to ATRX at the protein, RNA, or transcriptional level that impact *ATRX* function. There may be less frequent mutations in certain classes of genes that are responsible for creating more accessible chromatin, similarly to *ATRX*, or there may be other changes that occur via a different pathway that promote development of ALT. While we sought to identify ALT in a relatively large population, we found only very low levels of ALT in the non-pHGG examined and thus were unable to make observations regarding the impact of ALT on the clinical outcomes of these patients or the mutational landscape. Future work will focus on larger cohorts of non-pHGG to determine the true frequency of ALT in these patients and to determine the clinical significance of ALT in these groups.

Our analysis relied primarily on the presence of C-circles to identify ALT. However, in a small subset of tumors, ALT may occur without C-circles^44^. Additionally, due to inherent tumor and microenvironment heterogeneity, it is possible that some areas of the tumor may be truly ALT+, whereas the areas from which the DNA extracted would not have sufficient c-circles to register as positive^36^. By orthogonally validating our CCA with measurement of UBTF, we partially address this concern.

While our data do not show any difference in clinical survival in patients with pHGG with or without ALT, ALT+ cancers are often treatment-resistant cancers and novel therapies are needed for these patients ^3^. ALT-directed therapy remains an attractive target and may help sensitize ALT cancers to traditional cytotoxic chemotherapy^45^. Therapy that targets telomerase has been shown to induce ALT activity, which suggests an underlying intra-tumoral heterogeneity of telomerase lengthening mechanisms, and if ALT-directed therapy is developed clinically, it may be beneficial to pair this with telomerase-directed therapy^46^. Since ALT-directed therapy is not clinically available, it is not clear whether ALT-directed therapy could induce reactivation of telomerase. Current potential targets which might have therapeutic benefit in ALT+ tumors include ATM inhibitors, p53 reactivation, inhibitors that selectively target cells with *ATRX* mutations and G-quadruplex stabilizers^47^. Clinical trials exploring the dual use of PARP and ATR inhibitors relies on loss of function mutations in *ATRX* or *DAXX* to identify ALT+ patients48 are ongoing, thus developing a robust method to identify ALT in patients will be critical. By validating *TelomereHunter* with CCA, we have identified a computational tool that with additional verification and approval, may be clinically feasible to identify patients with ALT, which may be important as ALT-directed therapies become available.

We showed that *ATRX* loss or mutation only occurs in a subset of ALT+ pHGG, which has two important clinical implications. First, *ATRX*-directed therapies may not be effective for a subset of ALT+ tumors for which alternative therapies will be needed. Second, there are likely other major pathways driving the development of ALT in our patient population and loss of *ATRX* function, as measured by loss of ATRX protein expression or *ATRX* mutation, cannot be used alone as a metric for determining ALT status in patients.

By demonstrating an association between ALT and loss of MMR function in pediatric pHGG patients, we may have identified an area for potential therapeutic targets to disrupt the function of ALT and lead to tumor senescence, though this finding ought to be validated in larger cohorts as they become available. Our future work will continue to focus on identifying the key molecular changes that drive the development of ALT in pediatric brain tumor patients. We will validate the association between loss of MMR function and development of ALT, and work to understand whether loss of MMR function creates a permissive environment that promotes the development of ALT. Once this has been established, we will work to identify key targets that are essential for the ongoing ALT in these patients, with the goal of developing novel targeted therapies that disrupt ALT, sensitize cells to traditional cytotoxic chemotherapy and promote tumoral senescence. Our future work will additionally focus on elucidating the molecular differences in *ATRX* mutant versus *ATRX* wild-type ALT+ pHGG.

## Funding

This work was supported by NIH grant U2C-CA233285 (KAC), NIH 5T32CA009615-30 (JLS), NIH 2K12HD043245-16 (JLS), NIH R03-CA23036 (SJD), NIH Contract No. HHSN261200800001E (SJD), an Alex’s Lemonade Stand Foundation Young Investigator Award (JLR), the Matthew Larson Foundation (KAC), the Marlene Shlomchik Fellowship for Cancer Research (JLS), the Division of Neurosurgery at the Children’s Hospital of Philadelphia (PJS, ACR), and the Intramural Research Program of the Division of Cancer Epidemiology and Genetics of the National Cancer Institute.

## Conflicts of Interest

Dr. Angela J. Waanders is a member of the Scientific Advisory boards for Alexion and DayOne Biopharmaceuticals.

## Acknowledgements

We would like to thank the patients and families who have donated tissue for this research and we would like to acknowledge both the Children’s Brain Tumor Network and the Pacific Pediatric Neuro-Oncology Consortium for collecting, sequencing, and harmonizing the genomic and clinical data used for this study.

## Table Legends

**Table 1. Demographics and clinical information for patients with pHGG, separated by ALT status.** Patients with ALT+ pHGG are older and no differences in sex, race, ethnicity, tumor location or survival were noted. Mutations in *ATRX* were more common in ALT+ tumors and TMB is higher in ALT+ tumors. Germline variants in MMR genes are associated with ALT+ status (p=0.02).

**Table 2. Pediatric pHGG patients with pathogenic germline variants in MMR pathway genes have tumors enriched for ALT**

Listed are the seven patients from our primary analysis pHGG cohort (N = 85) and two patients from a clinical validation cohort in which we found a predicted pathogenic (P) or likely pathogenic (LP) germline variant in an MMR pathway gene. Somatic MMR and *ATRX* alterations predicted to be oncogenic are listed. *TelomereHunter* ratios, TMB, C-Circle, and UBTF are also shown. *Note: patient C641691 had a self-reported *PMS2* germline variant and was not included in statistical calculations.

**Figure S1.**
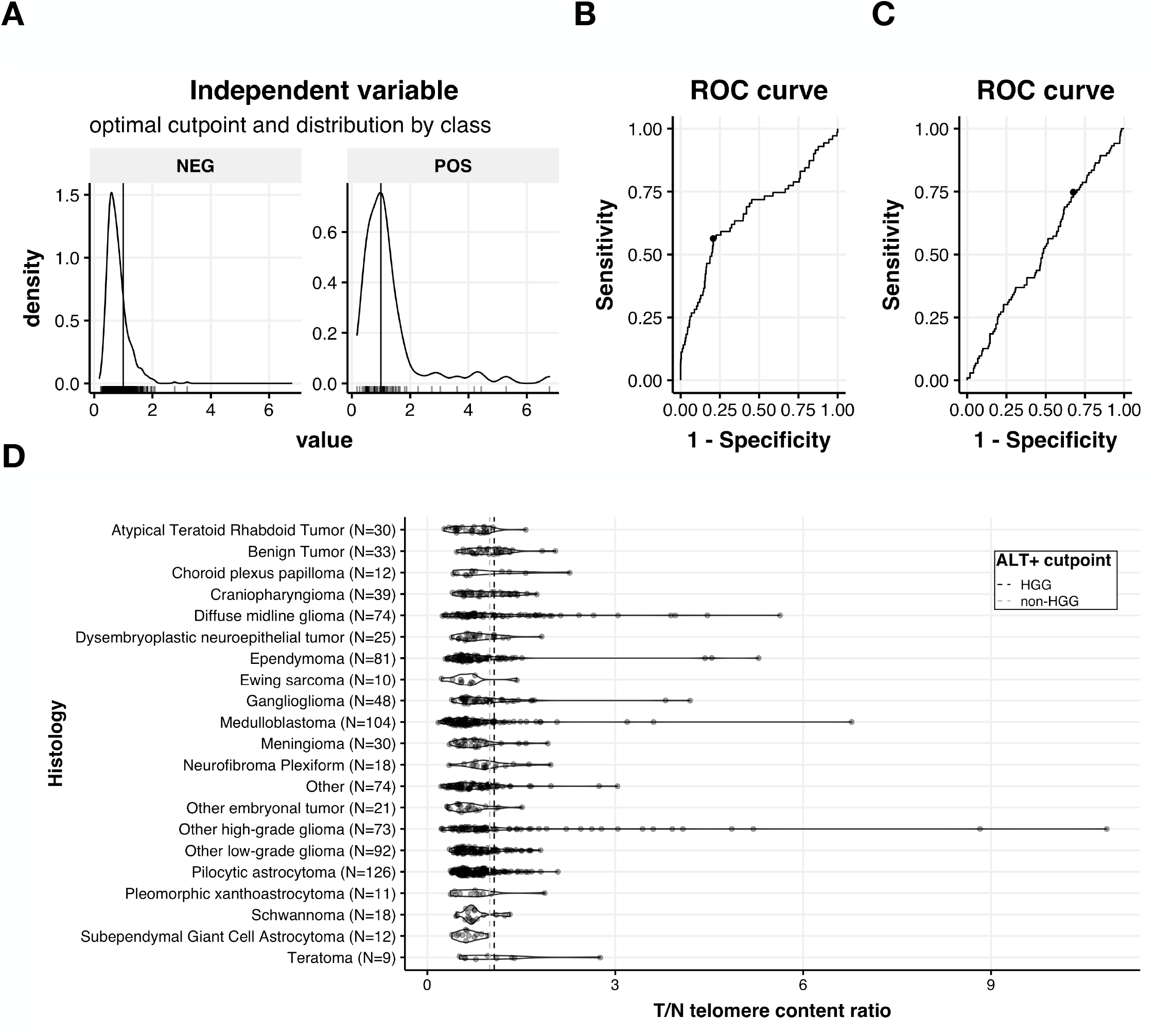
TelomereHunter performance on non-HGAT tumors in the PBTA. Using the C-circle assay data as the “truth” set for determine an ALT phenotype, we used the R package *cutpointr* to determine the optimal telomere ratio cutpoint for determining ALT +/-status. (**A**) Density plots for ALT + or - non-HGAT samples with a cutpoint of 0.9963 (x-intercept). (**B**) This cutpoint enabled an AUC = 0.66, 77% accuracy, 56.3% sensitivity, and 79.3% specificity, compared to shuffled telomere ratios (**C**), which resulted in an AUC = 0.526, 37.9% accuracy, 74.6% sensitivity, and 32.4% specificity. (**D**) Violin and sina plots of tumor/Normal (T/N) telomere content ratios across PBTA histologies. Cutpoints for ALT positivity are drawn as x-intercepts.

**Figure S2.**
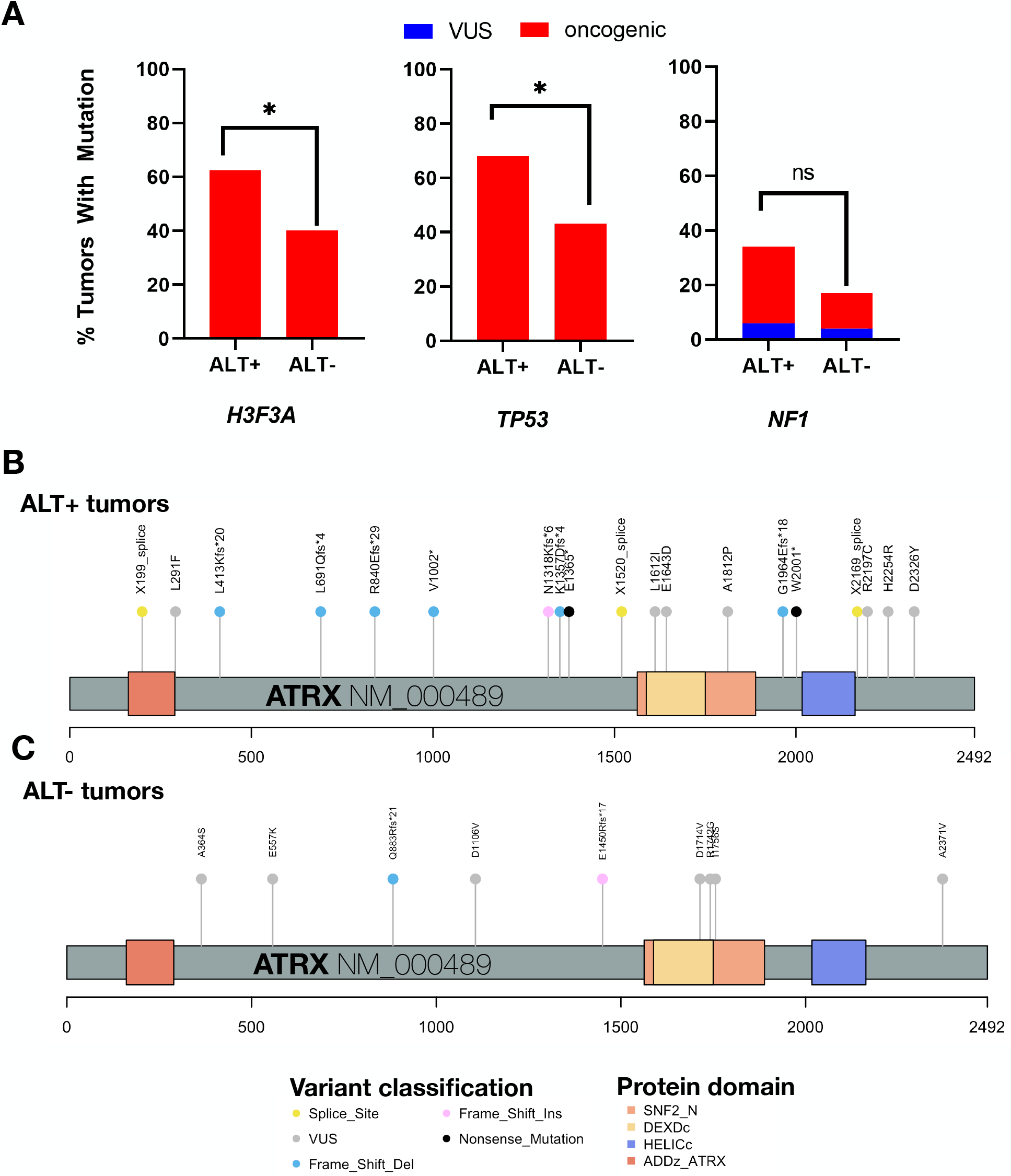
ALT+ HGGs have a higher prevalence of likely oncogenic ATRX mutations and higher TMB. (**A**) Barplots of ALT+/-tumors with somatic mutations in *H3F3A, TP53, or NF1*. Lollipop diagrams of protein changes due to *ATRX* mutations in ALT+ (**B**) or ALT-(**C**) HGG tumors. Colored variant classifications were all categorized as “likely oncogenic”, while those in grey were categorized as “VUS”, by oncoKB. The ATRX protein and its domains are also labeled. Of note, each mutation was unique to one sample.

**Figure S3.**
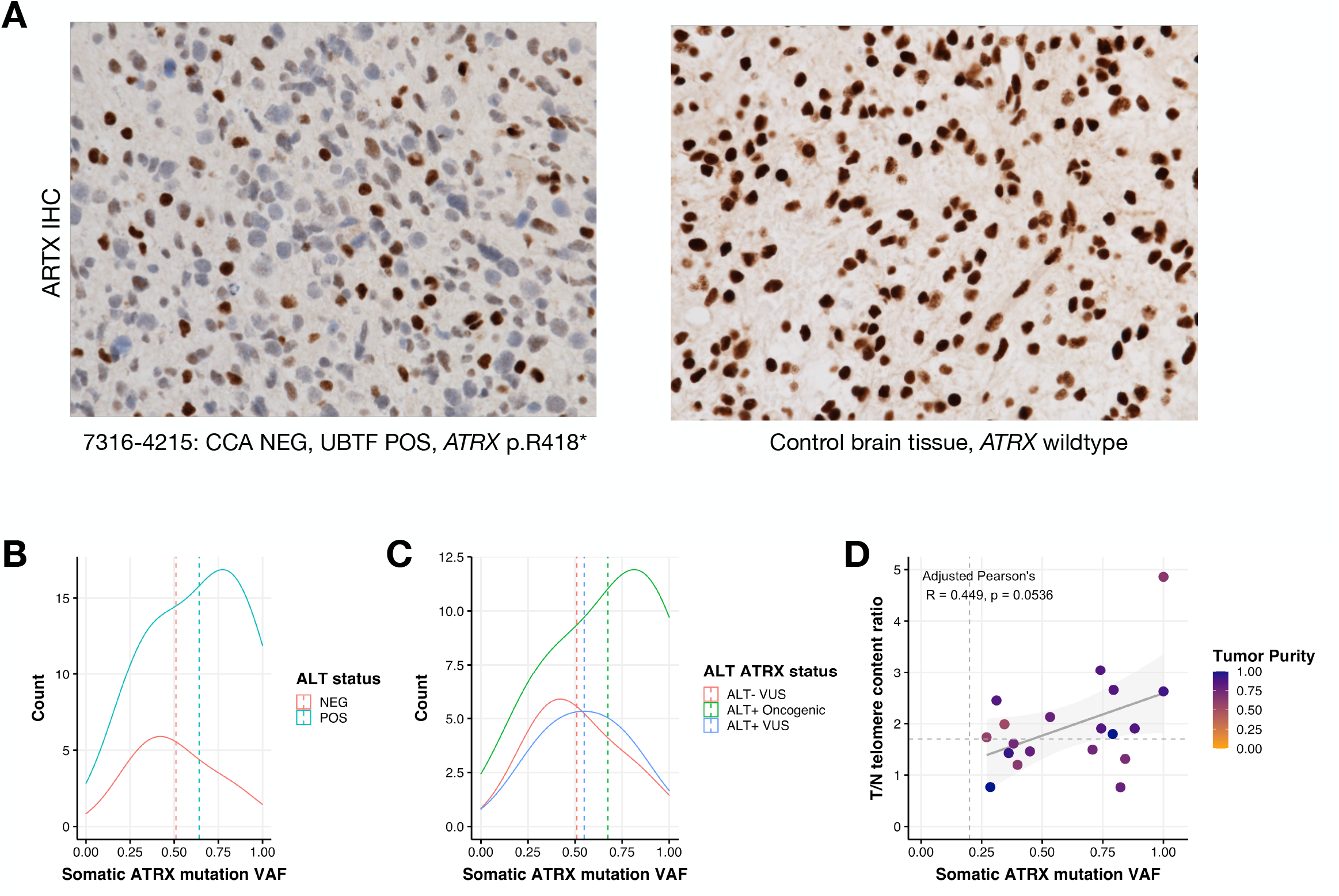
Intratumoral ATRX heterogeneity and clonality in pHGGs. (**A**) Left, ATRX IHC for *ATRX-*mutant tumor 7316-4215 showing intratumoral heterogeneity of ATRX. Tumor is negative for CCA and positive for UBTF. Right, IHC performed for an ATRX wild type control brain tissue. (**B**) density plot for somatic *ATRX* VAF by ALT status demonstrating and (**C**) ALT status and *ATRX* VAF by predicted oncogenicity of the mutation. Means of the VAF distributions are denoted by dotted lines at x-intercepts. (**D**) Pearson correlation of *ATRX* VAF by tumor/normal (T/N) telomere content ratio with points colored by tumor purity estimated by Theta2. All *ATRX* mutations are clonal (x-intercept dashed line denotes VAF = 0.20). All samples above the y-intercept dashed line (*TelomereHunter* cutoff = 1.699) are ALT+.

**Figure S4.**
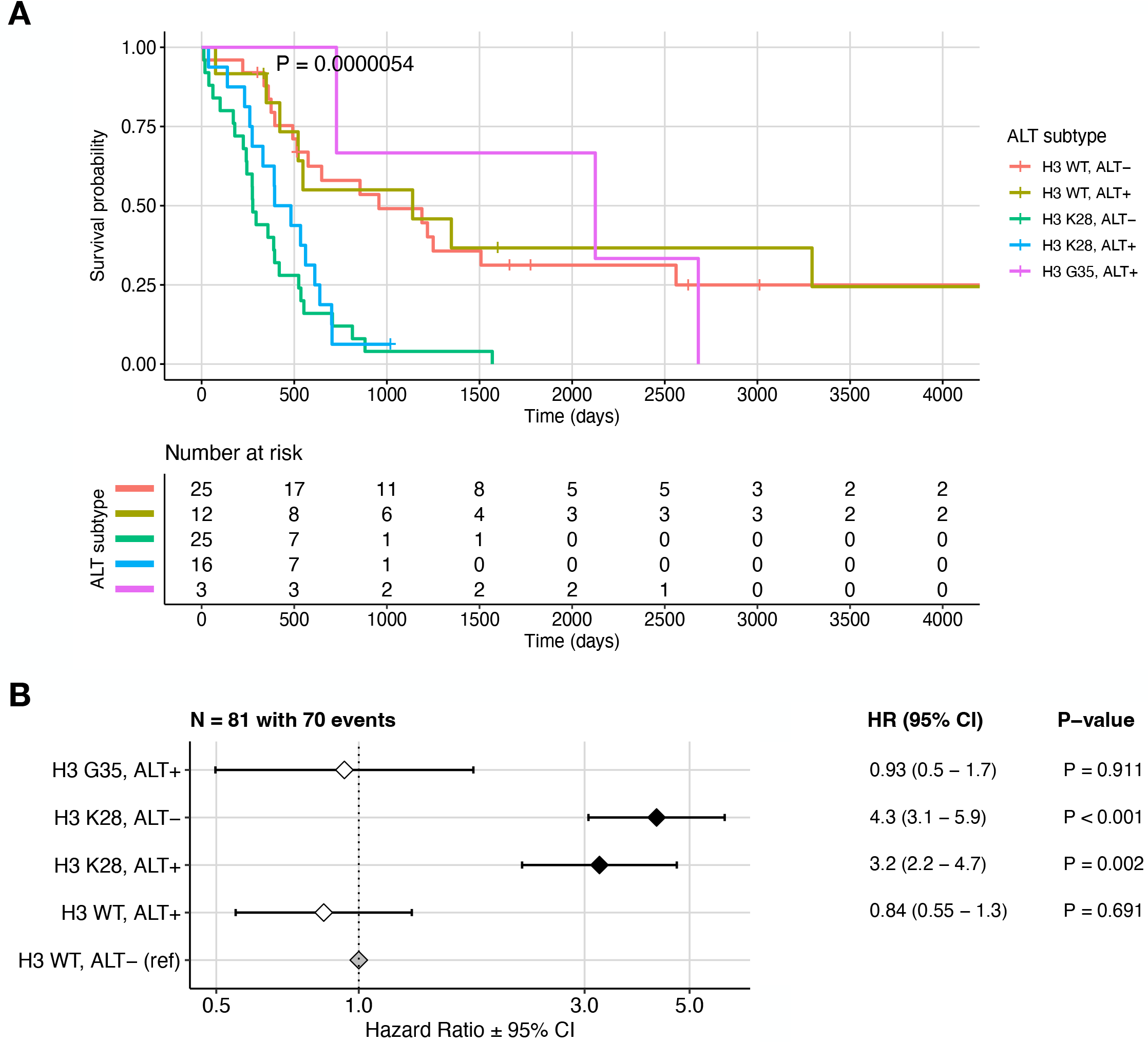
ALT positive H3 K28-altered DMGs tumors have a survival benefit. (A) There is a significant prognostic risk for H3 K28-altered HGATs, compared to H3 wild-type (WT) or H3 G35-mutant tumors (Kaplan-Meier log-rank test p = 5.4e-6). Multivariate cox regression was performed on the same subgroups. Hazard Ratios (HR), 95% confidence intervals (CI), and p-values are plotted in (B) versus the reference H3 WT, ALT-tumors.

## Notes

https://github.com/d3b-center/PBTA-ALT-analysis

### Summary of Updates

Addition of two authors and response to reviewer comments. Update allowed per policy at https://www.elsevier.com/about/policies/sharing

