## Supplementary_Legends for "ALT in Pediatric High-Grade Gliomas Can Occur without *ATRX* Mutation and is Enriched in Patients with Pathogenic Germline MMR Variants"

**Supplementary Figures and Tables**

**Figure S1. TelomereHunter performance on non-HGG tumors in the PBTA.** Using the C-circle assay data as the “truth” set for determine an ALT phenotype, we used the R package *cutpointr* to determine the optimal telomere ratio cutpoint for determining ALT +/- status. (**A**) Density plots for ALT + or - non-HGG samples with a cutpoint of 0.9963 (x-intercept). (**B**) This cutpoint enabled an AUC = 0.66, 77% accuracy, 56.3% sensitivity, and 79.3% specificity, compared to shuffled telomere ratios (**C**), which resulted in an AUC = 0.526, 37.9% accuracy, 74.6% sensitivity, and 32.4% specificity. (**D**) Violin and sina plots of tumor/Normal (T/N) telomere content ratios across PBTA histologies. Cutpoints for ALT positivity are drawn as x-intercepts.

**Figure S2. ALT+ pHGGs tumors have a higher prevalence of likely oncogenic *ATRX* mutations.**

(**A**) Bar plots of ALT+/- tumors with somatic mutations in *ATRX*, *TP53*, *NF1*, or *H3F3A*. Lollipop diagrams of protein changes due to ATRX mutations in ALT+ (**B**) or ALT- (**C**) pHGG tumors. Colored variant classifications were all categorized as “likely oncogenic” while those in grey were categorized as “VUS” by oncoKB. The ATRX protein and its domains are also labeled. Of note, each mutation was unique to one sample.

**Figure S3. Intratumoral ATRX heterogeneity and clonality in pHGGs.** (**A**) Left, ATRX IHC for *ATRX-*mutant tumor 7316-4215 showing intratumoral heterogeneity of ATRX. Tumor is negative for CCA and positive for UBTF. Right, IHC performed for an ATRX wild type control brain tissue. Images were taken at 40X. (**B**) density plot for somatic *ATRX* VAF by ALT status demonstrating and (**C**) ALT status and *ATRX* VAF by predicted oncogenicity of the mutation. Means of the VAF distributions are denoted by dotted lines at x-intercepts. (**D**) Pearson correlation of *ATRX* VAF by tumor/normal (T/N) telomere content ratio with points colored by tumor purity estimated by Theta2. All *ATRX* mutations are clonal (x-intercept dashed line denotes VAF = 0.20). All samples above the y-intercept dashed line (*TelomereHunter* cutoff = 1.068) are ALT+. Samples with TMB >=10 were excluded from analyses in B-D.

**Figure S4. ALT status alone is not a prognostic indicator for pHGGs.**

(**A**) There is a significant prognostic risk for H3 K28-mutant HGATs, compared to H3 wild-type (WT) or H3 G35-mutant tumors (Kaplan-Meier log-rank test p = 5.4e-6). Multivariate cox regression was performed on the same subgroups. Hazard Ratios (HR), 95% confidence intervals (CI), and p-values are plotted in (**B**) versus the reference H3 WT, ALT- tumors.

**Table S1. Tissue Microarray Data.** Patient and sample IDs are provided with results of the CCA, UBTF, ATRX IHC TMA. *ATRX* alterations, phase of therapy, and *TelomereHunter* ratios are also listed. From the primary analysis cohort (N = 85), 31 HGAT tumor samples were included in the TMA (N = 30) or CCA (N = 1). An additional 35 HGAT tumors (validation cohort) were analyzed for ATRX IHC (N = 35) or CCA (N = 2).

**Table S2. High Grade Astrocytic Tumor Table.** Full details are provided for all 85 unique patients analyzed in our study. Sample ID, *TelomereHunter* ratio, ultra-bright telomeric foci, ATRX, H3K28me3, and H3K28M IHC staining, CCA result, ALT designation, *ATRX* mutation status, *ATRX* mutation variant allele frequency (VAF), TMB, presence of germline and somatic MMR alterations are listed. Clinical and demographic information is provided, including CNS region of primary tumor, patient race, sex, and ethnicity, as well as integrated diagnosis, molecular subtype, age at diagnosis, age at last update and overall survival information is provided.

**Table S3. KEGG MMR gene list.** KEGG pathway MMR gene list used for germline pathogenicity analysis.

**Table S4. Survival Table.** Overall survival (OS) data is shown (first tab) for 85 patients in our primary analysis HGAT cohort. Kaplan-Meier log-rank results are shown in tab two with median OS and SD for each grouping of H3 subtype and ALT status. The third tab contains results of cox-multivariate analysis for the additive model “molecular subtype + ALT status”. The fourth tab contains all log-rank pair-wise comparisons with BH-corrected p-values.
