## Supplementary_Methods for "ALT in Pediatric High-Grade Gliomas Can Occur without *ATRX* Mutation and is Enriched in Patients with Pathogenic Germline MMR Variants"

***Tissue Microarray***

Construction of a pediatric brain tumor tissue microarray (TMA) was approved by the Children’s Hospital of Philadelphia Institutional IRB. Well-characterized HGAT patient formalin-fixed paraffin embedded tumors plus control tissues were punched in duplicated and sectioned into 5uM sections ^1^. Sections were stained by H&E, a routine glioma immunohistochemistry panel including ATRX, H3 K28M, and H3 K28me3, and presence of ultra-bright telomeric foci (UBTF), as described below.

***Measurement of Tel-FISH***

UBTF, which are markers of Alternative lengthening of Telomeres (ALT), were analyzed on a subset of patients on the pediatric brain tumor TMA described above. The IF-FISH protocol was performed as described previously ^2^. The C-rich/leading strand Telomere PNA probe (CCCTAAx3), labeled with Cy3 (PNA Bio #F1002) were used.

Stained slides were imaged using the Keyence BZ-X810 microscope at 4x magnification to create a map of the tissue section and to identify areas for high resolution imaging. Selected regions were then imaged at 40x magnification using filters for DAPI, Cy3, and Cy5. Cy3 channel images for the TelC-Cy3 probe were taken at a standard exposure time of 20ms. The Cy5 channel image exposure time for ATRX was 588ms. Multiple Z-stacks were taken for each imaging region at a Z-step of 1.0-3.0 μm. Image processing was performed using the Fiji program. IJ1 Macro scripts were written to convert 40x image tiles to 8-bit and then to stack their individual focus planes together. The resulting multi-stack image tiles for each fluorescence channel were merged to create a composite image tile. These composites were then stitched together using the Grid/Collection stitching plugin in Fiji ^3^. For images captured using the Keyence microscope, tiles are imaged in a snake-by-rows order with approximately 30% overlap between adjacent tiles. The stitching function was also set to compute tile overlap to simultaneously register adjacent images and thus more accurately avoid the presence of overlapping regions. The stitched Cy3 channel image was then multiplied by 15 for better staining visualization. The resultant image was qualitatively analyzed for the presence of ultra-bright telomeric foci (extremely bright intracellular signals of varying size and intensity despite a very short exposure).

***Immunohistochemistry***

α-ATRX antibody rabbit polyAb (Sigma Aldrich HPA001906), α-Histone H3 K27M antibody rabbit mAb (Histone H3, Sigma Aldrich SAB5600095), and α-Tri-methyl-Histone H3 (Lys27) (C36B11) rabbit mAB (Cell Signaling #9733) immunohistochemistry was used to stain 5 uM formalin fixed paraffin embedded tissue microarray (TMA) sections. Staining was performed with primary antibody incubation performed for one hour at room temperature at 1:1000 (ATRX), 1:1000 (H3K28M), or 1:150 (H3K28me3) dilution. Antigen retrieval, dehydration and imaging was performed as described previously^4^.

***Pediatric brain tumor data and genomic analyses***

*Somatic mutation and CNV calling*

Somatic mutations were called using Strelka2, Mutect2, Lancet, and VarDict as described by the OpenPBTA project ^5^. Consensus mutations were retained if they were called by two or more of the four variant callers and augmented with hotspot mutations if found by only one of the four methods, as described within the OpenPedCan project (<https://github.com/PediatricOpenTargets/OpenPedCan-analysis>, release v10). CNVs were called using ControlFREEC, CNVkit, and MantaSV with consensus being a CNV present in two of the three methods. This method and further annotation of CNVs and tumor purity is described in detail in the OpenPBTA ^5^. The oncoprint was generated using the R package ComplexHeatmap ^6^.

*Tumor Mutation Burden and ATRX, H3F3A, TP53, and NF1 mutation analysis*

Tumor mutation burden (TMB) was obtained from the OpenPBTA project (<https://github.com/AlexsLemonade/OpenPBTA-analysis>, release-v22-20220505) and defined as the number of consensus SNVs per effectively surveyed coding sequence of the genome. OncoKB ^7^ was used to annotate somatic mutations as “Oncogenic”, “Likely oncogenic” or “Unknown” (variants of unknown significance, VUS). Intronic, silent, and RNA variants were removed. Using the maftools R package ^8^, lollipop plots of the ATRX protein were created separately for ALT+ and ALT- HGATs. The OpenPedCan v10 data was used to assess whether ALT+ or ALT- HGATs contained mutually exclusive or co-occurring somatic mutations.

*Survival analysis*

Tumors were annotated by H3 gene mutation status (molecular subtype) and ALT status, as determined by C-circle, if available, or *TelomereHunter.* Log-rank analysis was performed by adapting methods from the OpenPBTA, and Kaplan-Meier was plotted. An additive multivariate cox regression analysis was performed for “H3 molecular subtype+ALT status”, followed by log-rank pairwise comparisons between all groups, with p-values corrected by Benjamini-Hochberg.

*Code availability*

Code to reproduce the above somatic analyses is available at <https://github.com/d3b-center/PBTA-ALT-analysis>.
